# Pharmacological inhibition of HDAC6 downregulates TGF-β via Smad2/3 acetylation and improves dystrophin-deficient muscles

**DOI:** 10.1101/2022.01.21.477182

**Authors:** Alexis Osseni, Aymeric Ravel-Chapuis, Isabella Scionti, Yann-Gaël Gangloff, Vincent Moncollin, Remi Mounier, Pascal Leblanc, Bernard J. Jasmin, Laurent Schaeffer

## Abstract

The absence of dystrophin in Duchenne muscular dystrophy (DMD) disrupts the dystrophin dystroglycan glycoprotein complex (DGC) resulting in fibers fragility and atrophy, associated with fibrosis and microtubules and neuromuscular junction (NMJ) disorganization. The specific non-conventional cytoplasmic histone deacetylase 6 (HDAC6) was previously shown to regulate acetylcholine receptor distribution and muscle atrophy. Here we show that administration of the HDAC6 specific inhibitor tubastatin A to the DMD mouse model *mdx* improves muscle strength, restores microtubules, NMJ and DGC organization, and reduces muscle atrophy and fibrosis. These effects involve the known action of HDAC6 on microtubules acetylation and muscle atrophy but also involve a yet undiscovered action of HDAC6 on transforming growth factor beta (TGF-β) signaling. Conversely, to inhibitors of nuclear HDACs that regulate TGF-β signaling *via* the activation of Follistatin expression, HDAC6 inhibition acts downstream of TGF-β ligands and receptors by increasing Smad2/3 acetylation in the cytoplasm which in turn inhibits its phosphorylation and transcriptional activity.

---

Duchenne muscular dystrophy (DMD) is an X-linked neuromuscular recessive disorder affecting around 1 in 3.500 newborn males worldwide and is the most common and fatal form of muscular dystrophy^1,2^. Patients with DMD manifest their first clinical symptoms at the age of 3-4 years and become wheelchair dependent between the ages of 7 and 13 years. The ambulation period can be prolonged in many boys with DMD with early initiation of steroid treatment. The terminal stage of the disease starts when patients require assisted ventilation by the age of around 20 and patients usually die in their third or fourth decade due to respiratory or cardiac failure^3–7^. DMD results from mutations in the dystrophin gene that cause the synthesis of nonfunctional or the absence of dystrophin protein. Dystrophin is a critical component of the dystrophin-associated glycoprotein complex (DGC) in muscle^8^. DGC is a structure that spans the sarcolemma and forms a mechanical link between the cytoskeleton and the extracellular matrix *via* the association of dystrophin with both actin and microtubule cytoskeleton and the binding of dystroglycan to laminin in the basal lamina, respectively^9,10^. During muscle contraction, the DGC acts as a molecular shock absorber and stabilizes the plasma membrane^11,12^. Loss of dystrophin is associated with muscle deterioration and degeneration and prevents the DGC to exert its functions, thus rendering muscle fibers more susceptible to contraction-induced membrane damage leading to cell death^13–16^. This pathologic process is accompanied by inflammation and fibrosis that participates to muscle wasting and loss of function^17–21^.

Despite tremendous research efforts, no cure is available for DMD patients yet. Gene-based therapeutic strategies, such as exon skipping, suppression of stop codons, Adeno-associated virus (AAV)-mediated mini-dystrophin delivery, or CRISPR/Cas9 gene editing are actively being investigated to treat DMD^22,23^. In parallel, pharmacological treatments are also being developed^24–27^. Such approaches act on specific signaling pathways and cellular events including those that can cause upregulation of utrophin A^28–30^. Nonetheless, glucocorticoids still serve as the gold standard therapy, acting mostly as anti-inflammatory drugs^31^. With the use of steroids and multidisciplinary care, particularly mechanical ventilation, lifespan expectancy of DMD patients has been considerably increased and affected individuals can now reach 30 to 40 years of age.

Pan-deacetylase inhibitors were previously shown to significantly improve function and morphology in dystrophin-deficient mice^32,33^. Indeed, deacetylase inhibitors conferred dystrophic muscles resistance to contraction-coupled degeneration and alle*via*ted both morphological and functional consequences of the primary genetic defect. The mechanism involved was shown to be the activation of the expression of the activin binding protein follistatin that sequesters many ligands of transforming growth factor beta (TGF-β) receptors, thereby inhibiting this TGF-β pathway. TGF-β signaling plays a central role in promoting muscle atrophy and fibrosis in neuromuscular disorders. A variety of ligands including GDF8/myostatin, GDF11 and Activins bind to type II TGF-β receptors that trigger the phosphorylation of Smad2 and Smad3 transcription factors upon activation. Smad2/3 phosphorylation enables oligomerization with Smad4 and translocation in the nucleus to activate the expression of genes involved in muscle atrophy in cooperation with FoxO transcription factors^34–36^.

As compared to other nuclear Histone Deacetylases (HDACs) known, HDAC6 is described as a unique cytoplasmic member of the HDAC family, belonging more specifically to class IIb^37,38^. HDAC6 deacetylates many cytoplasmic substrates such as tubulin, cortactin and HSP90 whereas other HDACs are typically central regulators of gene expression. Indeed, classical HDACs such as HDAC4 and HDAC5, are involved in the regulation of atrogenes *via* Myogenin and Foxo3 transcription factors^39,40^. In this context, we have previously shown that HDAC6 is an atrogene activated by FoxO3a that can interact with the ubiquitin ligase atrogin1/MAFbx^41^. Considerable efforts to develop specific HDAC6 inhibitors have been made to treat a variety of diseases. In particular, several anti-cancer treatments targeting HDAC6 have been proposed^42,43^. Furthermore, HDAC6 inhibition has been shown to be beneficial for some neurodegenerative diseases, including Amyotrophic lateral sclerosis or Charcot-Marie-Tooth disease^44,45^. The HDAC inhibitor, tubastatin A (TubA) stands out as a highly specific HDAC6 inhibitor. Indeed, TubA efficiently and specifically inhibits HDAC6 deacetylase activity with an IC50 of 15 nM and a strict selectivity for HDAC6 over all other HDACs (over 1000-fold), except for HDAC8 (57-fold)^45–47^. TubA is known to inhibit TGF-β-induced type-1 collagen expression in lung fibroblasts^48,49^ and previous reports have shown that TGF-β increases the activity of HDAC6^48,50^. Indeed, Smad7 regulates expression of HDAC6 in prostate cancer in response to TGF-β^51^ and HDAC6 may play an indispensable role in balancing the maintenance and activation of primordial follicles through mechanistic target of rapamycin (mTOR) signaling in mice^52,53^.

Previous works with pan-HDAC inhibitors have shown that HDAC inhibition improves the dystrophic phenotype^27,33^. However, it remains unclear which HDAC is involved in this beneficial effect. In a recent study, we discovered that HDAC6 regulates microtubule stability and acetylcholine receptors (AChR) clustering in muscle fibers^54^. We thus wondered whether HDAC6 could play a role in the patho-mechanism of DMD and be used as a novel therapeutic target for the disease. To test this, we investigated the effect of TubA, a highly specific inhibitor of HDAC6, in DMD mouse model *mdx* and C2C12 cells. Altogether, our results show that the pharmacological inhibition of HDAC6 deacetylase activity with TubA is beneficial to muscle from *mdx* mice, both at the functional and morphological levels. Investigation of the mode of action of TubA revealed that HDAC6 inhibition down-regulates TGF-*β* signaling through an increase of Smad2/3 acetylation in skeletal muscle.

## Results

To evaluate the effect of the inhibition of HDAC6 deacetylase activity in dystrophic-deficient mice, 7-week-old *mdx* mice received daily intraperitoneal injections of TubA (*mdx*-TubA) or vehicle (*mdx*-veh) at a dose of 25 mg/kg/day as previously described^45,55^. The mice were treated for 30 days to allow comparison with the previous pre-clinical studies using this duration of treatment in *mdx* mice^56–58^. Untreated wild-type mice (WT-CTL) mice were used as baseline controls (Fig. 1a). The efficiency of the treatment with TubA was first evaluated by measuring the increase in tubulin acetylation (Fig. 1b). As shown in Fig. 1c and 1d, TubA treatment for 30 consecutive days caused as expected, a large increase in α-tubulin acetylation in TA muscle from *mdx* mice. In line with published data^54,55^, quantification of the relative level of acetylated tubulin showed a 36.5% ± 7.8% increase following treatments with the specific HDAC6 inhibitor (P < 0.001 compared with *mdx*-veh mice). To confirm that HDAC6 inhibition did not affect histone acetylation, histone H3 acetylation on lysine 9 (ac-H3K9) and total histone H3 levels in TA muscles were evaluated by Western blot. Consistent with ongoing muscle damage in *mdx* mice, H3K9 acetylation was increased in TA muscles from *mdx* mice compared to WT-CTL animals (+43.8% ± 14.6%; extended data Fig. 1a, b). As expected, TubA did not increase H3K9 acetylation in *mdx* mice (P > 0.05). Together, these data indicate that intraperitoneal injection of TubA efficiently and specifically inhibits HDAC6 deacetylase activity in muscle.

**Fig. 1.**
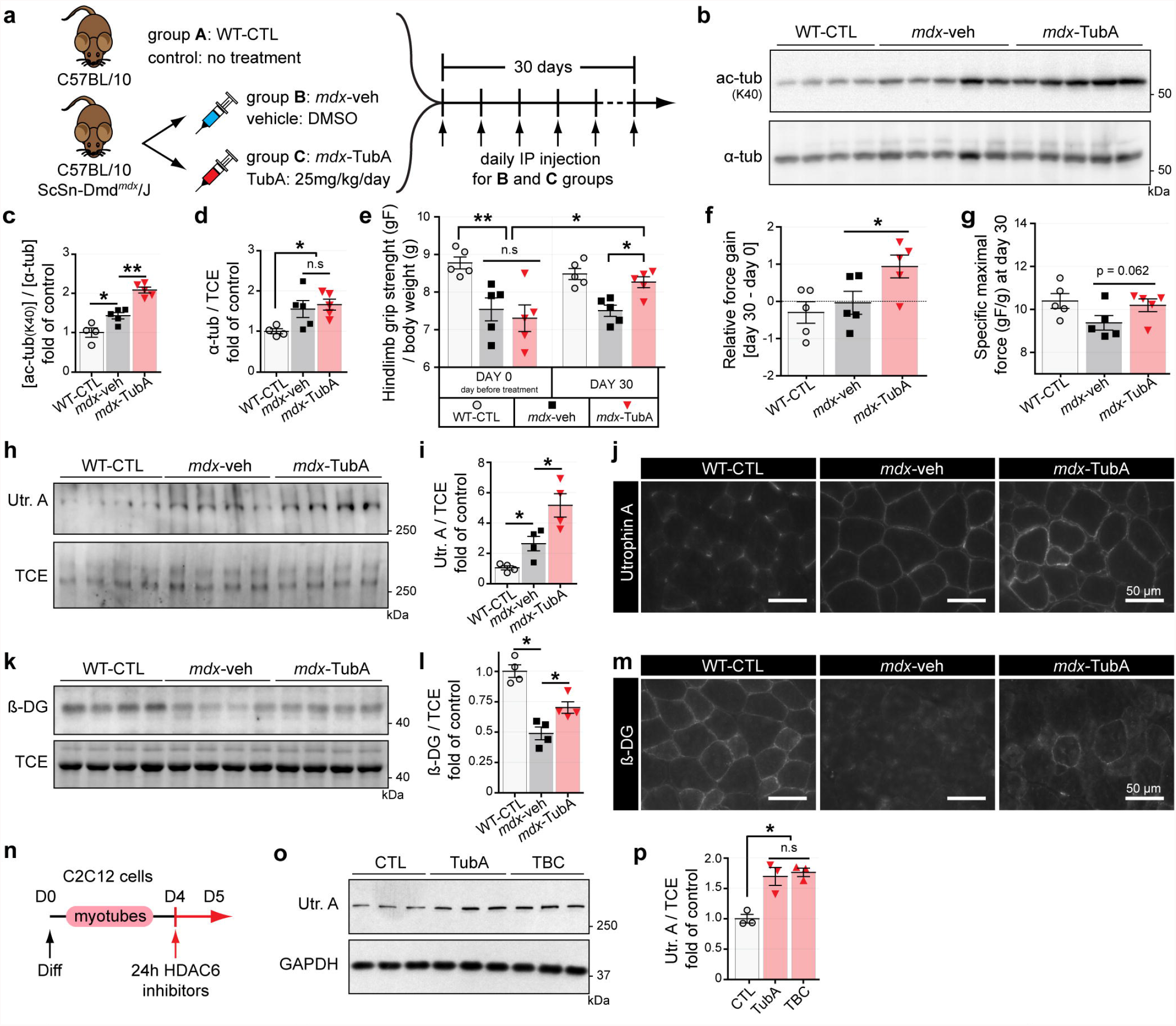
*In vivo* in *mdx* mice, HDAC6 inhibition via TubA treatment increases grip strength, sarcolemmal localization of utrophin A and promotes reassembly of the DGC. **a**, Protocol of TubA treatment. Three group of 7-wk-old mice have been evaluated for 4 weeks either without treatment (group A, C57BL/10 mice: WT-CTL), or treated with daily injection for 30 consecutive days with DMSO (group B, C57BL/10ScSn-Dmd*mdx*/J mice; *mdx*-veh) or with DMSO supplemented with TubA at 25 mg/kg/day (group C, C57BL/10ScSn-Dmd*mdx*/J mice; *mdx*-TubA). **b**, To evaluate the level of tubulin acetylation (ac-tubK40) in TA muscles, Western blot analysis were performed. **c, d**, Quanti fications of acetylated tubulin **(c)** and α-tubulin (α-tub, **d)** protein levels were respectively normalized to α-tubulin and TCE (4-5 mice per group). **e**, Grip strength was measured on a grid measuring maximal hindlimb grip strength normalized on body weight (5 mice per group). **f**, Relative force gain was calculated by the difference between grip strength measured at the last day (day 30) and the day before starting treatment (day 0). **g**, Specific maximal force was evaluated by the best score of grip strength obtain in each animals at day 30. **h, l, k, l**, To evaluate levels of Utrophin A (Utr. A) and β-dystroglycan (β-DG) in TA muscles, Western blot analysis **(h**,**k)** and quantification (**i,l**) were performed. TCE was used as a loading control (4-5 mice per group). **j, m**, Cross sections of TA muscle from WT-CTL, *mdx*-veh or *mdx-TubA* were stained with an antibody against Utrophin A (**j**, in gray) or against β-dystroglycan(**m**, in gray). Scale bars: 50 µm. **n**, 4-d-old C2C12 myotubes pretreated for 24 h with different HDAC6 inhibitors TubA (5 µM) and tubacin (TBC, 5 µM) or with DMSO (CTL, 1 µl). **o**, Representative Western blots showing Utrophin A. GAPDH was used as a loading control. **p**, Quantification of Utrophin A protein levels normalized with GAPDH (3 independent experiments quantified). Quantifications show means ± SEM. *, P < 0.05; **, P < 0.01; n.s, not significant, P > 0.05; Mann-Whitney U test. kDa, relative molecular weight in kiloDalton.

Next, we evaluated the effect of the TubA treatment on muscle strength in *mdx* mice. Seven-week-old untreated *mdx* mice exhibited significantly lower grip strength for all paws (P < 0.01) compared with WT-CTL mice. After 30 days of treatment with TubA, the grip strength of *mdx* mice was significantly increased by 1.5-fold (P <0.05 compared with *mdx*-veh mice; Fig. 1e), with a substantial force gain over 30 day (P <0.05; Fig. 1f). TubA treatment almost completely restored muscle strength of *mdx* mice to WT levels (P = 0.5476 compared with WT-CTL mice; Fig. 1e). Interestingly, the maximal force was also increased by 35% ± 9.0% in TubA-treated *mdx* mice compared to *mdx*-veh animal although it did not reach the significance level due to interindividual variability (P = 0.062; Fig. 1g). Altogether, these data indicate that inhibition of HDAC6 deacetylase activity in *mdx* mice restores muscle strength to WT levels.

Because of its functional and structural similarity with dystrophin, Utrophin A can compensate for the lack of dystrophin in DMD^24,29,59,60^. In adult and healthy muscle, Utrophin A is exclusively localized at synapses^61,62^ and it was shown that Utrophin A expression all along muscle fibers can efficiently compensate for the lack of dystrophin^24,29,63–65^. *Mdx* mice present an overall milder phenotype than DMD patients^66^. This is partially explained by the compensatory up-regulation of sarcolemmal Utrophin A expression^24,25,29,59,60,67,68^. All groups of animals described in figure 1a were thus analyzed by Western blot and immunofluorescence to evaluate the levels of Utrophin A. As expected, Utrophin A levels were increased at the sarcolemma (P < 0.05, Fig. 1h) in *mdx* mice compared with WT-CTL mice. TubA-treatment further increased Utrophin A levels by ∼2-fold compared with *mdx*-veh mice (P < 0.05; Fig. 1i). Immunofluorescence experiments further established that TubA treatment indeed caused an increase in sarcolemmal Utrophin A levels in *mdx* mouse muscles, thereby possibly conferring a higher protective effect on muscle fiber integrity (Fig. 1j). Additionally, we assessed both the amount and localization of β-dystroglycan (β-DG), a member of the DGC, in order to determine whether TubA treatment caused reassembly of the DGC along the sarcolemma. As expected, in the absence of dystrophin, β-DG accumulation at the plasma membrane is strongly decreased (Fig. 1k, m). Western blot quantification revealed that TubA increased the amount of β-DG in *mdx* mice by 43.2% ± 7.0% (P < 0.05; Fig. 1l). Immunofluorescence experiments with TubA (Fig. 1m) indicated that in addition to the increase in Utrophin A levels, β-DG accumulation at the plasma membrane was increased, suggesting that TubA treatment restored the DGC complex.

To determine whether the effects of TubA on DGC expression was also observed in cultured muscle cells, C2C12 myotubes were treated for 24 hours with 5µM of one of two specific inhibitors of HDAC6 deacetylase activity: TubA and tubacin^69^ (TBC; Fig. 1n). After the treatment, Utrophin A expression was evaluated on whole cell extracts by Western blot (Fig. 1o). Both HDAC6 inhibitors induced a significant upregulation of Utrophin A levels ∼1.75-fold (P < 0.05) in C2C12 muscle cells compared to vehicle-treated cells (Fig. 1p). Together, these data demonstrate that treatments with HDAC6 inhibitors increase Utrophin A levels in C2C12 myotubes in agreement with the data obtained with the DMD mouse model *mdx*.

To determine whether TubA treatment provided additional benefits to *mdx* muscle fibers, muscle atrophy was evaluated. Histopathological studies have shown that abnormal fiber size distribution, with a strong increase in very small fibers, is a hallmark of dystrophic muscles^70^. Fiber size distribution was assessed by measuring the cross-sectional area (CSA) of individual fibers in the slow oxidative *soleus* muscle (SOL) and the fast glycolytic *extensor digitorum longus muscle* (EDL), from *mdx* mice treated with TubA or vehicle and control mice. As expected, the CSA profiles of vehicle-treated *mdx* muscles displayed substantial heterogeneity and many small fibers when compared to healthy muscles (Fig. 2a, d). In all muscles analyzed, TubA restored the fiber size distribution comparable to that in WT muscles by significantly decreasing the proportion of small muscle fibers (Fig. 2b, e). In TubA-treated *mdx* mice, these muscle adaptations were accompanied by a significant decrease of the average variance coefficient. Indeed, TubA treatment reduced by ∼ 25% (P < 0.01, compared with *mdx*-veh mice) the variance coefficient of fibers CSA in *mdx* EDL and SOL muscles (Fig. 2c, f). In summary, four weeks of TubA treatment normalized fiber size distribution in both EDL and SOL *mdx* muscles indicating a protective effect of TubA against muscle atrophy.

**Fig. 2.**
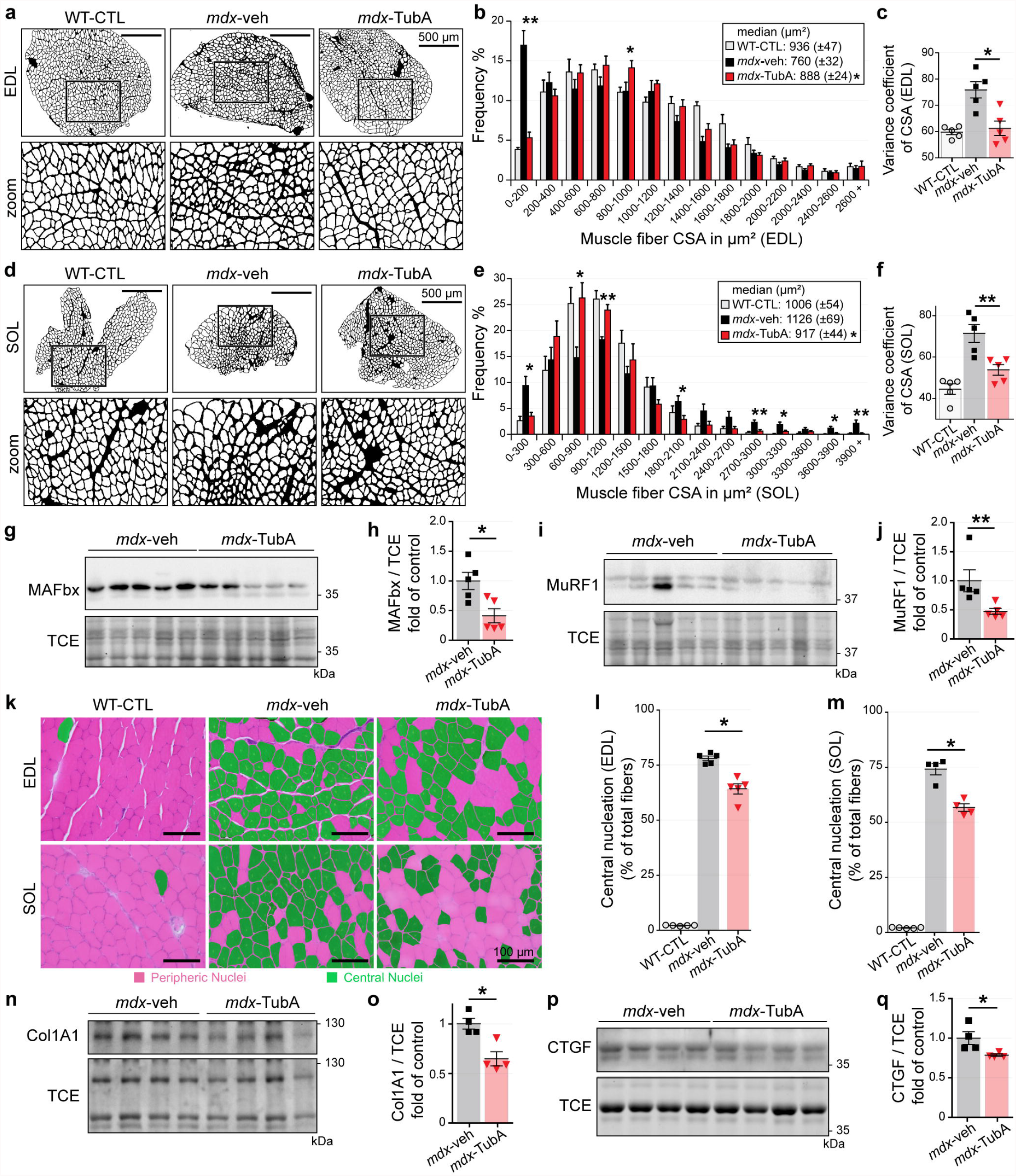
TubA treatment improves and restores DMD phenotype in *mdx* muscle and protects from atrophy. **a, d**, Cross-section areas (CSA) of entire EDL (a) and SOL **(d)** muscles from 11-wk-old C57BL/10 mice (WT-CTL) and C57BL/10ScSn-Dmd*mdx*/J mice treated with TubA (*mdx*-TubA) or with vehicle-OMSO (*mdx*-veh) for 30 consecutive days were stained using laminin staining and then binarized on lmageJ. Scale bars: 500 µm. **b, e**, Graphical summary of CSA (4-5 mice per group). Quantifications show means± SEM. *, P < 0.05; **P < 0.01; two-way ANOVA (*mdx*-TubA versus *mdx*-veh). Median CSAs of each muscle are displayed above the frequency histograms. **c, f**, Measure ments of variance coefficient in EDL (**c**) and SOL **(f)** muscle fibers (4-5 mice per group). **g, h, i, j**, To evaluate levels of MAFbx **(g, h)** and MuRF1 (**i, j**) in TA muscles, Western blot analysis **(g, i)** and quantifications (**h, j**) were performed (5 mice per group). **k**, Representative examples of cross-sections of EDL and SOL muscles were stained using hematoxylin and eosin. Scale bars: 100 µm. Centrally nucleated fibers are colored in green. **l, m**, Percentage of central nucleation in EDL (**l**) and SOL **(m)** muscle fibers (4-5 mice per group). **n, o, p, q**, To evaluate levels of collagen type I alpha 1 (**n, o**, Col1A1) and connective tissue growth factor (**p, q**, CTGF) in TA muscles, Western blot analysis **(n, p)** and quantifications **(o, q)** were performed (4 mice per group). TCE was used as a loading control for all Western blots. Quantifications show means± SEM. *, P < 0.05; **P < 0.01; n.s, not significant, P > 0.05; Mann-Whitney U test. kDa, relative molecular weight in kiloDalton.

We previously showed that HDAC6 contributes to muscle atrophy^41^. Given the overall increase in fiber size observed in *mdx* muscles upon TubA treatment, we investigated the mechanism by which pharmacological inhibition of HDAC6 decreased muscle atrophy in *mdx* mice (Fig. 2g, i). The expression of protein markers involved in muscle atrophy was evaluated by Western blot. TubA caused a 57% ± 11% reduction of MAFbx/atrogin1 (P < 0.05; Fig. 2h) and a 35% ± 4% reduction of the E3 ubiquitin-protein ligase TRIM63/MuRF1 (P < 0.05; Fig. 2j) compared with *mdx*-veh animals. Together, these data indicate that HDAC6 inhibition reduces expression of key mediators of muscle atrophy, thus providing an explanation for the normalization of the size of muscle fibers.

In addition, heterogeneity in *mdx* fiber size results mainly from the presence of small regenerating fibers subsequent to the loss of necrotic fibers. Central nucleation is as a key indicator of muscle damage in dystrophic muscle fibers^70,71^. To evaluate the effect of TubA on the extent of muscle damage, the number of fibers with central nuclei was evaluated using hematoxylin eosin staining (Fig. 2k). TubA caused a significant decrease in the number of centronucleated fibers (−15% ± 2% in EDL muscle, Fig. 2l and - 24 % ± 2% in SOL muscle, Fig. 2m; both P < 0.05, compared with *mdx*-veh mice). Furthermore, the loss of muscle fibers is accompanied by the progressive accumulation of fibrotic tissue^72^. We therefore investigated by Western blot if the expression of fibrosis-associated proteins in *mdx* muscles was reduced by TubA (Fig. 2m, p). The protein levels type I Collagen Alpha 1 Chain (Col1A1) and connective tissue growth factor (CTGF) were reduced in *mdx* mice treated with TubA compared to vehicle treated *mdx* mice (−36% ± 7%, Fig. 2o and – 16% ± 1%, Fig. 2q; respectively, P < 0.05). Together, these data show that HDAC6 inhibition in dystrophic muscle reduces central nucleation, normalizes fiber size distribution, reduces the proportion of small fibers, and diminishes fibrosis.

Dystrophin and Utrophin A link the DGC complex to the microtubule network^10,73^. In agreement with previous data^10,74^, we confirmed by Western blotting and immunofluorescence staining that *mdx* mice have more tubulin and a greater microtubule density, respectively, in TA and EDL fibers *mdx* mice than WT-CTL mice (P < 0.05, Fig. 1d and extended data Fig. 2a, b). However, TubA treatment did not affect overall α-tubulin abundance (P > 0.05 compared with *mdx*-veh mice; Fig. 1d). In healthy muscle, the microtubule network forms a grid lattice with longitudinal, transverse, and perinuclear microtubules^75–77^. In healthy skeletal muscle, transverse and longitudinal microtubules are regularly spaced by ∼2 µm and ∼5 µm, respectively (see arrowheads Fig. 3a and extended data Fig. 2c-e). In *mdx* muscles, immunofluorescence experiments show a disorganization of the microtubule network, with a loss of the grid-like organization (see arrows Fig. 3a). Interestingly, in *mdx* mice treated with TubA the microtubule lattice was restored (see the spacing between microtubules on the blue and yellow line scans and extended data Fig. 2c-e). We further analyzed the microtubule network with a software specifically developed to analyze microtubule directionality^78^ (TeDT program; Fig. 3b). TeDT program analysis revealed a significant decrease of transversely oriented microtubules (centered around 90°) in vehicle-treated *mdx* muscles compared to WT and to TubA-treated *mdx* mice (P < 0.05 compared with *mdx*-veh mice). These data show that TubA treatment restores the organization of the microtubule network in *mdx* muscles. In agreement with previous work^54,79^, this further demonstrate that microtubule acetylation *via* HDAC6 inhibition stabilizes and restores transverse microtubules.

**Fig. 3.**
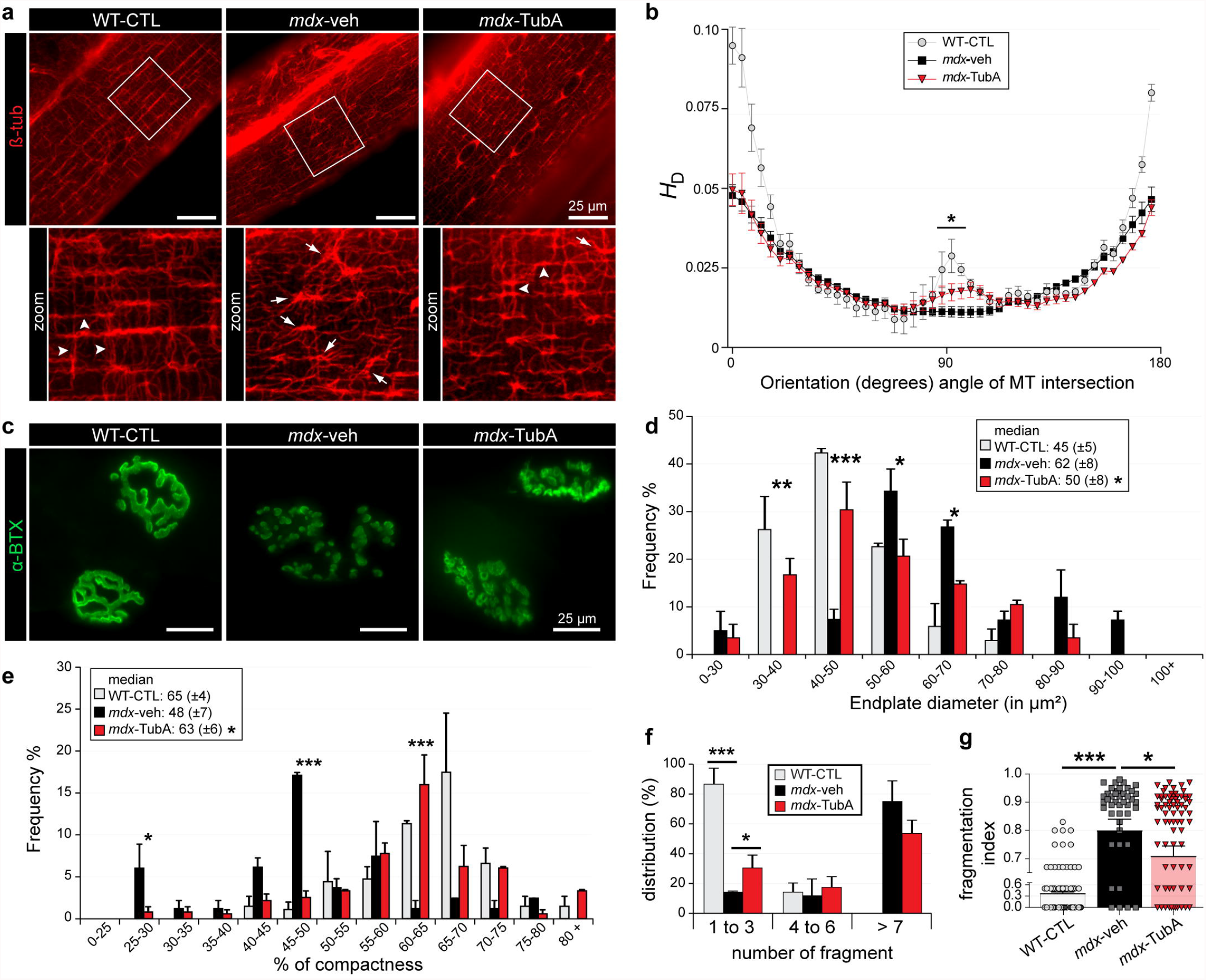
TubA treatment stabilizes MT network and protects NMJ morphological characteristics from dystrophic mice. **a**,**c**, Isolated fibers of TA from 11-wk-old C57BL/10 mice (WT-CTL) and C57BL/1 0ScSn-Dmd*mdx*/J mice treated with TubA (***mdx***-TubA) or with vehicle-OMSO (*mdx*-veh) for 30 consecutive were stained with an antibody against β-tubulin to label MT network (**a**, in red) or stained with α-BTX-A488 (**c**, in green) to label NMJs. Scale bars: 25 µm. **b**, MT network organization analysis of four to six fibers from two or three mice using TeDT software. The final generated graph presents a global score for each given degree of MT orientation, with 0 and 180 degrees corresponding to the longitudinal MT and 90 degrees corresponding to the transverse MT Arrowheads represent regular organization of MT network whereas arrows show disorganization of the MT network, with a loss of the grid-like organization. **d, e**, Graphical summary of NMJ endplate diameter **(d)** and NMJ compactness **(e)**. Median CSAs of each muscle are displayed above the frequency histograms (40-75 NMJs counted). **f, g**, Distribution of number of fragments **(f)** and Fragmentation index **(g)** have been quantified (43-73 of NMJs counted). Quantifications show means ± SEM. *, P < 0.05; **, P < 0.01; ***, P < 0.001; (*mdx*-TubA versus *mdx*-veh), Mann-Whitney U test.

Describe to ameliorate the microtubule organization, HDAC6 also regulates AChR clustering at the NMJ^54^. The severe fragmentation of NMJs observed in mdx muscles strongly suggests that dystrophin participates to the maintenance of the NMJ^80^. We therefore analyzed NMJ organization in control and treated *mdx* mice (Fig. 3c). *α*-bungarotoxin staining used to visualize the acetylcholine receptor indicates that NMJ organization was severely compromised in control *mdx*-veh treated mice as compared to control WT NMJs (Fig. 3c, d), as expected from previous works^81–83^. Based on a fragmentation index and the number of clusters per NMJ and the endplate diameter, AChRs in *mdx* mice are more discontinuous and punctate than those in WT mice. The compactness of *mdx*-veh NMJs was decreased by 26% ± 11% (P < 0.05; Fig. 3e) and the fragmentation index was increased by approximately 2-fold (P < 0.001; Fig. 3f, g). Interestingly, TubA-treated *mdx* NMJs compared to vehicle-treated *mdx* NMJs showed a normalization of endplate diameter (−24% ± 13%, P < 0.05; Fig. 3d) and of endplate compactness (+ 31.3% ± 12.5%, P < 0.05; Fig. 3e). The number of fragments composing the NMJ was also increased (+ 63% ± 1% of NMJs composed of 1 to 3 fragments, P < 0.05; Fig. 3f) and the fragmentation index was decreased by 13.2% ± 2% (P < 0.05; Fig. 3g). Altogether, TubA-treatment partially rescued AChR distribution in *mdx* mice thanks to an increase of compactness and recruitment of AChR patches.

Furthermore, dystrophin transcription is also controlled via mTOR pathway^84^. Indeed, mTOR deficiency leads to reduced muscle dystrophin content and causes dystrophic defects leading to severe myopathy. In this context, we investigated if HDAC6 inhibition in *mdx* muscle could promote protein synthesis by measuring the phosphorylation of various components of the mTOR pathway (Fig. 4a). mTOR protein level was not significantly changes (P = 0.342; Fig. 4b, c) whereas a ∼2-fold increase of p70-S6 kinase phosphorylation at threonine 389 was observed (P < 0.05; Fig. 4d, e). Consistent with increased p70-S6 kinase activity, we observed a ∼2-fold increase of the phosphorylation of S6 on both Ser235-236 and Ser240-244 (P < 0.05; Fig. 4f, g, h). Finally, TubA induced a significant increase in the phosphorylation of the translation inhibitor 4EBP1 on the mTOR-sensitive sites as shown by the increase in the ratio of 4E-BP1 γ over 4E-BP1 β isoforms for phospho-Thr36-45, phosphor-Thr70 as well as for total 4E-BP1 (Fig. 4i, j, k, l). Altogether, our data show that a 30 day-treatment with TubA stimulates muscle protein synthesis in *mdx* mice *via* the mTOR pathway.

**Fig. 4.**
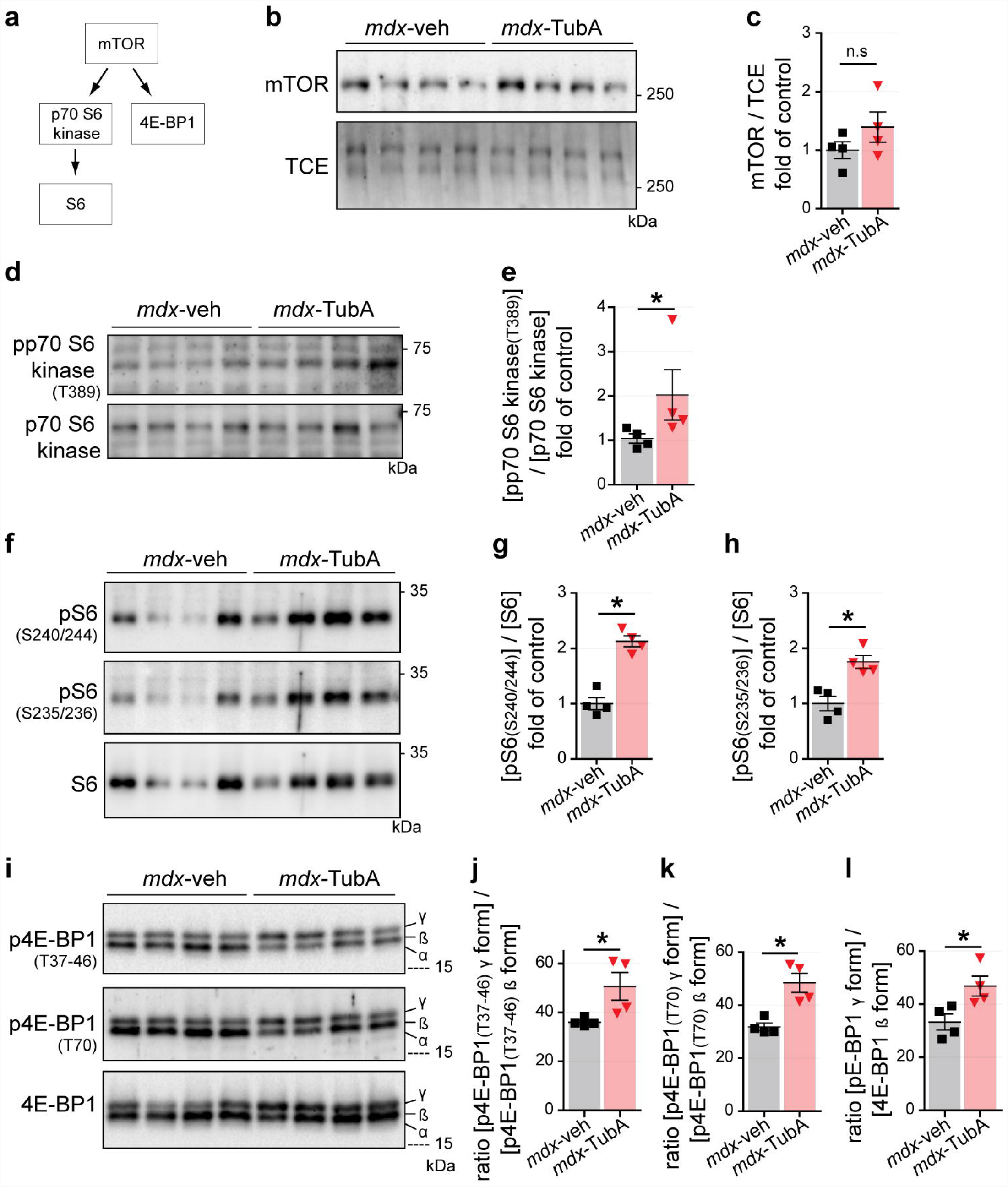
HDAC6 inhibition actives mTOR pathway in *mdx* mice. **a**, Schematic summary of downstream targets of mTOR pathway. 11-wk-old C57BL/10ScSn-Dmd*mdx*/J mice treated with TubA (*mdx*-TubA) or with vehicle-DMSO (*mdx*-veh) for 30 consecutive days have been analyzed in TA muscle by Western blot analysis **(b, d, f, i)** and quantifications of mTOR **(c)**, pp70 S6 kinase (**e**, T389), pS6 (**g**, S240-244), pS6 **(h**, S235/236), p4E-BP1 (**j, T**37/46), p4E-BP1 (**k**, T70), and 4E-BP1 **(l)** were been performed (4 mice per group). TCE was used as a loading control. Quantifications show means ± SEM. *, P < 0.05; n.s, not significant, P > 0.05; Mann-Whitney U test. kDa, relative molecular weight in kiloDalton.

Recently data reported that mTOR activation is led by the TGF-β signaling^85^ of which family members are major regulators of muscle mass. The effects of TubA thus prompted us to explore TGF-β signaling. Indeed, muscle atrophy induced by the TGF-β member myostatin involves Smad2/3 signaling that activates MAFbx expression and downregulates mTOR signaling^86,87^. Smad2/3 proteins require phosphorylation to enter inside nucleus and activate its target genes^88^. Interestingly, Smad2/3 proteins have also been shown to be acetylated in the nucleus^89,90^. Mouse C2C12 myoblasts were treated for 24 hours with either vehicle (DMSO), TubA or SB431542 (SB43) a specific inhibitor of the TGF-β/Activin/NODAL pathway^91^. After removal of the inhibitors, C2C12 myoblasts were exposed for 30 min to recombinant human TGF-β1^92^ (rhTGF-β1) and the subcellular localization of Smad2/3 was analyzed by immunofluorescence using anti-acetylated tubulin and anti-Smad2/3 antibodies.

TubA treatment similarly increased tubulin acetylation both in the presence and absence of rhTGF-β1, indicating that TGF-β signaling does not affect HDAC6 deacetylase activity (Fig. 5a and extended data Fig. 3a,c). In agreement with previous data^93,94^, rhTGF-β1 treatment increased Smad2/3 phosphorylation at serines 465-467 and 423-425, respectively (extended data Fig. 3b). Consistently, immunofluorescence and Western blot experiments in C2C12 cell line respectively indicated that rhTGF-β1 increased phosphorylation and nuclear accumulation of Smad2/3 (Fig. 5a, b and extended data Fig. 3a, b c), which were blocked by SB43 (Fig. 5a, b and extended data Fig. 3a, b). Similarly, Smad2/3 phosphorylation and nuclear accumulation were strongly diminished in the presence of TubA (Fig. 5c, e and extended data Fig. 3d, e), and this reduction correlated with an hyperacetylation of Smad3 (P < 0.05, Fig. 5d). TGF-β signaling is known to be upregulated in dystrophic muscles^95^. To confirm *in vivo* that HDAC6 inhibition downregulated TGF-β signaling, Smad2/3 acetylation and phosphorylation were evaluated by Western blot in TA muscles of *mdx* mice treated with TubA or DMSO (Fig. 5f, h). In TubA-treated *mdx* mice, Smad2/3 acetylation on Lys19 was increased by 74% ± 24% compared to DMSO-treated *mdx* mice (P < 0.05; Fig. 5g). This increase in acetylation was paralleled by a 55% ± 4% decrease of Smad3 phosphorylation on Ser432-425 (P < 0.05; Fig. 5i).

**Fig. 5.**
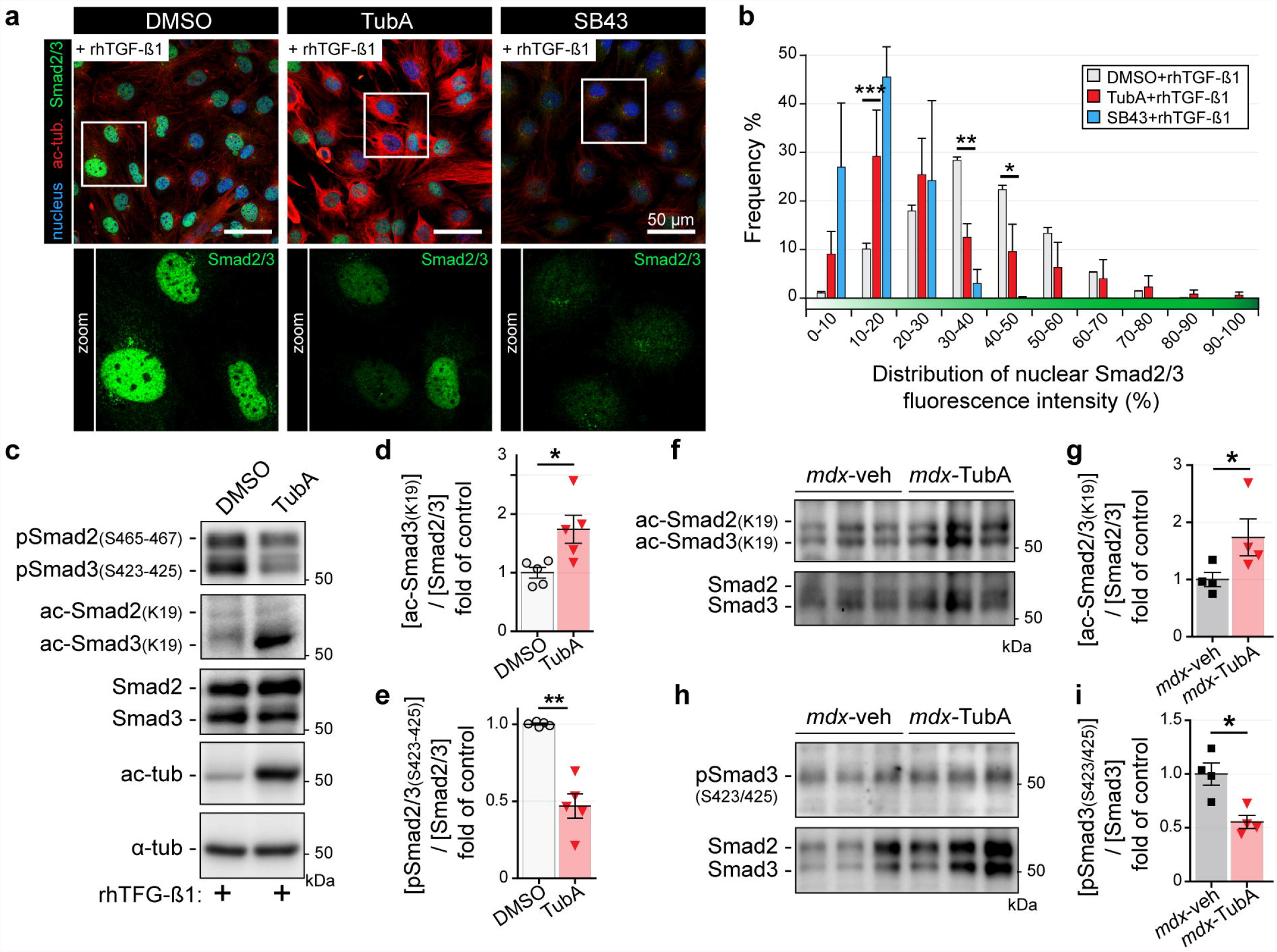
TGF-β signaling is regulated by TubA *via* acetylation of Smad2/3. **a-e**, 4-d-old C2C12 myoblasts pretreated for 24 h with either HDAC6 inhibitor(TubA, 5 µM), or selective inhibitor ofTGF-β1(S843, 5 µM) or with DMSO(CTL, 1 µI). Myoblasts were then treated for 30 min with rhTGF-β1(10 ng/ml). 11-wk-old C57BL/10ScSn-Dmd*mdx*/J mice treated with TubA(*mdx*-TubA) or with vehicle-DMSO (*mdx*-veh) for 30 consecutive days have been analyzed in TA muscle by Western blot analysis **(f, h)** and quantified **(g, i). a**, Myoblasts were double-stained with antibodies against Smad2/3(in green) and acetylated tubulin(ac-tub, in red). Nuclei were labeled with DAPI(in blue). **b**, Graphical summary of nuclear distribution of Smad2/3 fluorescence intensity from 3 independent experiments. Quantifications show means ± SEM. *, P < 0.05; **P < 0.01; two-way ANOVA (TubA+rhTGF-β1 versus DMSO+rhTGF-β1). **c, f, h**, Levels of Smad2/3 phosphorylation (S465-467; S423-425), Smad2/3 acetylation (K19), Smad2/3, acetylated tubulin(ac-tub), and α-tubulin(α-tub) were visualized by Western blot analysis. **d, e, g, i**, To evaluate levels of Smad2/3 acetylation **(d, g)** and Smad2/3 phosphorylations (**e, i**, both in C2C12 cells (**d, e;** 5 independent experiments quantified) and in TA muscles (**g, i**, 4 mice per group), quantifications have been performed. TCE was used as a loading control for all Western blots. Quantifications show means± SEM. *, P < 0.05; **, P < 0.01; Mann-Whitney U test. kDa, relative molecular weight in kiloDalton.

## Discussion

Here, we have evaluated the effect of HDAC6 inhibition by TubA in *mdx* mice and in C2C12 muscle cells. TubA treatment significantly ameliorated *mdx* muscle function and decreased the overall histopathological dystrophic features. In search of the mechanism of action of HDAC6, we discovered that HDAC6 regulated TGF-β signaling *via* the acetylation of Smad2/3, thus identifying Smad2/3 as new targets of HDAC6. TubA increased Smad2/3 acetylation, preventing Smad2/3 phosphorylation and nuclear translocation, thereby decreasing the expression of atrogenes such as MAFbx and MuRF1 and upregulating the mTOR pathway. Altogether, this mechanism can explain the beneficial effect of HDAC6 inhibition to counter muscle atrophy and fibrosis in *mdx* muscles (Fig.6).

**Fig. 6.**
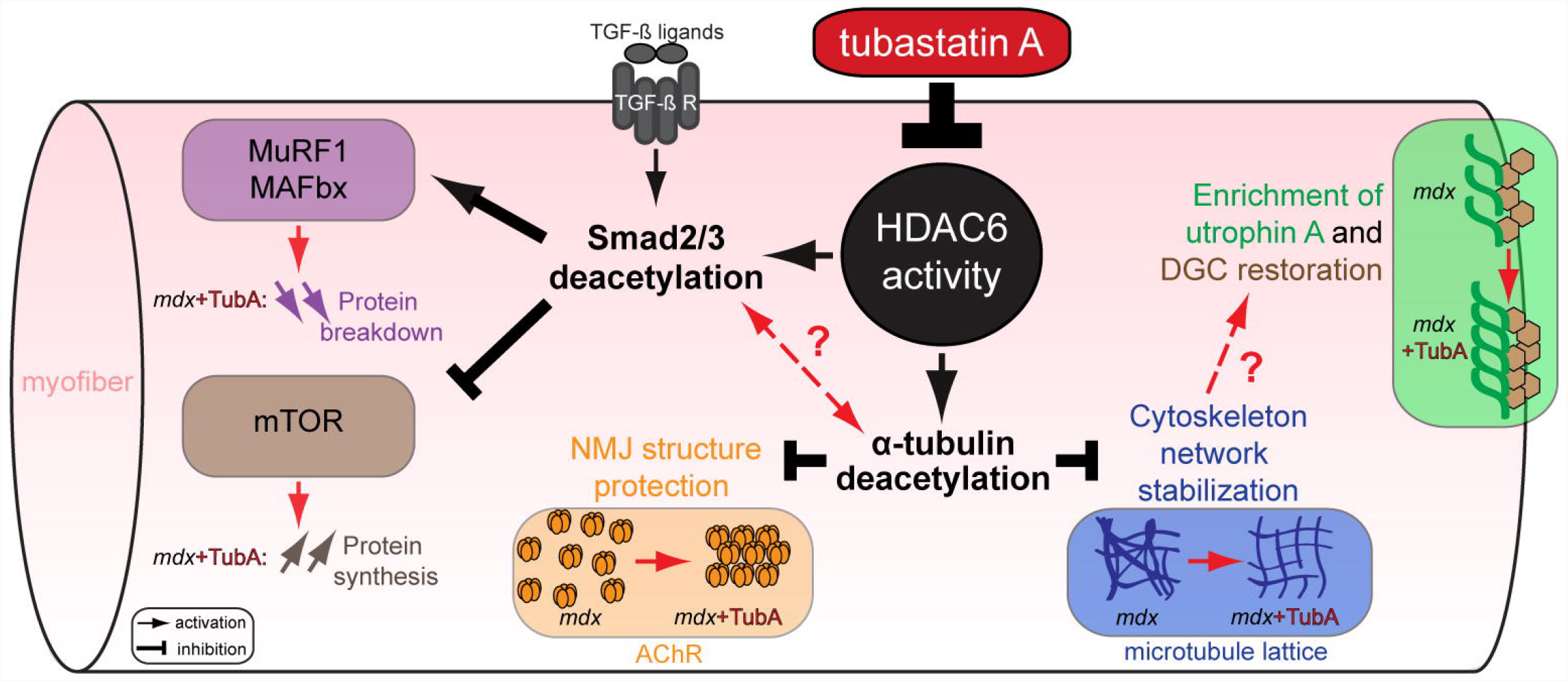
Consequences of HDAC6 activity inhibition regulation through TubA treatment in DMD mice model. HDAC6 inhibitor such as TubA induces a decrease in HDAC6 activity that leads to an acetylation of α-tubulin and of Smad2/3. HDAC6 pharmacological inhibition allows an increase of α-tubulin acetylation to restore DGC and stabilize MT network/NMJ organizations. Additionally, specific inhibition of HDAC6 increase acetylation of Smad2/3 which can interfere with TGF-β signaling to both reduce muscle atrophy by reducing the expression of key actors such as MAFbx/MuRF1 and to stimulate protein synthesis *via* mTOR pathway. Our results identify HDAC6 as a pharmacological target of interest for DMD.

Our results indicate that stabilizing microtubules by preventing their deacetylation *via* HDAC6 restores normal microtubule organization in dystrophin-deficient muscles. In these *mdx* muscles, the loss of force production is commonly attributed to structural impairments of the muscle fiber cytoskeleton and changes in signaling^96^. Interestingly, stabilization of the microtubule network was shown to protect against contraction-induced injury, suggesting that targeting the microtubule cytoskeleton may provide novel opportunities for therapeutic intervention in DMD^74^. Accordingly, *mdx* muscles treated with TubA contain less centronucleated fibers, possibly because muscle fibers are more resistant.

Our data show that TubA partially restores NMJ morphology in *mdx* mice. In DMD patients and *mdx* mice, NMJs are noticeably disorganized and have been shown to be associated with deficits in neuromuscular function / transmission, thus highlighting the contribution of NMJ impairment in altered function and recovery of dystrophic muscles^80,97–99^. We recently showed that HDAC6 is involved in microtubules organization at the NMJ, by regulating AChR distribution and NMJ organization^54^. In *mdx* mice, structural changes in microtubules at the NMJ are probably involved in the disorganization of the NMJ^82^. Our present study indicates that and protecting NMJ structure by stabilizing microtubules probably participates to the beneficial effect of TubA treatment on *mdx* muscle function. In addition, a significant increase of Utrophin A levels correlates with improved prognosis in DMD patients^100^. Therefore, the ability of HDAC6 inhibition to increase extra-synaptic Utrophin A levels probably participates to the preservation of *mdx* muscle integrity. How HDAC6 regulates Utrophin remains to be determined. A plausible hypothesis would be that the restoration of the microtubule network improves Utrophin trafficking to the membrane.

In contrast to most HDACs that directly act on gene expression *via* transcription regulatory complexes, HDAC6 is strictly cytoplasmic and none of its known substrates are transcription factors. Here we show that HDAC6 deacetylates Smad2/3 and regulates its nuclear accumulation. Smad2/3 proteins mediate the action of TGF-β signaling to promote protein catabolism and fibrosis and to inhibit protein anabolism. Inhibition of Smad2/3 nuclear accumulation by HDAC6 inhibition can therefore explain the beneficial effect of TubA on atrophy reduction and protein synthesis stimulation *via* the mTOR pathway. Beneficial effects of HDAC inhibitors such as Givinostat were previously demonstrated in *mdx* mice and DMD patients that stimulate the expression of the activin binding protein follistatin, whose main activity is to block TGF-β signaling^27,32,33^. These beneficial effects of HDAC6 on *mdx* muscles recapitulate effects of Givinostat. Moreover, in *mdx* and DMD patient muscles, inhibition of TGF-β activity attenuates both degeneration and fibro-calcification in *mdx* muscle^101^ and fibrosis^27^. Consistently, HDAC6 inhibition also reduced fibrosis in *mdx* muscles.

Interestingly, rather than directly affecting the expression of individual genes as shown for other HDACs, HDAC6 acts on TGF-β signaling by targeting Smad2/3 proteins in the cytoplasm before their translocation into the nucleus. Hence, HDAC6 regulates the downstream targets of Smad2/3, thus providing a novel pharmacological entry to interfere with TGF-β signaling. Smad2 and Smad3 share 92% sequence identity but their functions are not completely redundant. Smad2 knock-out mice die at embryonic day 10.5 with vascular and cranial abnormalities and impaired left-right patterning^102,103^ whereas Smad3 knock-out mice are *via*ble but suffer from impaired immune function and chronic inflammation^104,105^. In addition, they exhibit preferences for association with specific transcription factors. For example, Smad3 interacts preferentially with FoxO over Smad2^34^.. Interestingly, acetylation events reported for Smad2/3 this far are performed by nuclear factors^89,90^. Consistently most of these events have been shown to regulate promoter binding, transactivation activity, nuclear export or protein stability. In the nucleus, Smad2 and Smad3 are both acetylated at lysine 19 by p300/CBP in response to TGF-β, which increases their DNA binding activity and binding to target genes promoters^106^. Acetylation of Smad2 and Smad3 may selectively up- or down-regulate the expression of TGF-β-regulated genes. Of note, Smad2 and Smad3 can also be acetylated on several other lysines^89,90^.HDAC6 being strictly cytoplasmic, Smad deacetylation by HDAC6 probably affects different processes. Our results indicate that increasing Smad2/3 acetylation reduces their phosphorylation and then nuclear accumulation. We can thus speculate that preventing Smad2/3 deacetylation in the cytoplasm reduces their phosphorylation by TGF-β receptors and/or reduce their association with Smad4. Altogether, Smad2/3 acetylation in the nucleus is required to activate the expression of their target genes, whereas of Smad2/3 acetylation in the cytoplasm inhibits their function. Intriguingly, HDAC6 inhibition in C2C12 cells affects only Smad3 acetylation but both Smad2 and 3 phosphorylation. In muscle treated with TubA, the acetylation of both Smad2 and 3 was reduced. This could reflect and intrinsic difference between cultured muscle cells and muscle fibers or it could just be due to the fact that durations of *in vitro* and *in vivo* treatments were very different.

It is conceivable that after its activation by p300, acetylated Smad2/3 molecules that are exported back from the nucleus to the cytoplasm require deacetylation by HDAC6 to allow their re-entry in the TGF-β pathway. Therefore, HDAC6 inhibition would prevent Smad2/3 deacetylation in the cytoplasm thereby reducing the amount of Smad2/3 available to participate to TGF-β signaling and increasing the amount of cytoplasmic Smad2/3. Another possibility would be that Smad2/3 deacetylation by HDAC6 regulates its ubiquitination and subsequent degradation. Both hypothesis are consistent with the fact that HDAC6 inhibition increases the amount of cytoplasmic Smad2/3 protein.

In a previous work, we showed that HDAC6 expression was increased during muscle atrophy and that it participated to muscle wasting *via* a direct interaction with the ubiquitin ligase MAFbx^41^. Therefore, we propose that HDAC6 inhibition prevents the degradation of muscle proteins at two levels: inhibition of TGF-β signaling *via* Smad2/3 and inhibition of ubiquitination and subsequent degradation of MAFbx substrates. The inhibition of TGF-β signaling was also shown to be beneficial to limit cachexia and cancer progression in mouse models^107,108^. Many tumors secrete TGF-β and the finding that HDAC6 regulates TGF-β might also be relevant to explain the beneficial effects observed on cancer progression with HDAC6 inhibitors^51,52,109^.

Altogether our results show that HDAC6 pharmacological inhibition significantly ameliorates dystrophic dystrophic-deficient muscles *via* three complementary mechanisms (Fig. 6): i) microtubule stabilization that favors NMJ and DGC restoration, ii) increase in Utrophin A levels, iii) inhibition of TGF-β signaling via Smad2/3 targeting to reduce muscle atrophy and fibrosis and to stimulate protein synthesis.

## Supporting information

Extented Data 1

Extented Data 3

## Material and Methods

### Ethics statement, Animal models, treatments, and preparation of samples

All procedures using animals were approved by the Institutional ethics committee and followed the guidelines of the National Research Council Guide for the care and use of laboratory animals, and by University of Ottawa Animal Care Committee. All procedures were in accordance with the Canadian Council of Animal Care Guidelines. For *in vivo* experiments, both control C57 black 10 (C57BL10) and C57BL/10ScSn-Dmd*mdx*/J *mdx* were used (The Jackson laboratory Bar Harbor, USA). *Mdx* mice were treated for 30 consecutive days with either tubastatin A (TubA, APExBIO, #A4101; 25 mg/kg/day, intraperitoneally) solubilized in 2% DMSO or saline supplemented with 2% DMSO (vehicle-control)^45^. After treatment, muscles were dissected and either: i) frozen and crushed in liquid nitrogen for protein and RNA extraction or ii) embedded in Tissue-Tek OCT compound (VWR, Mississauga, Canada) and frozen in isopentane cooled with liquid nitrogen for cryostat sectioning^54^ or iii) were manually dissociated^77^ with the following modifications: single TA fibers were fixed 30 min at room temperature in PBS-4% paraformaldehyde, permeabilized 60 min in PBS-1% Triton X-100 at 30°C before saturation and incubation with antibodies as described below.

### Hindlimb grip strength

All injections and behavioral tests were performed in a blinded manner at the University of Ottawa Animal Behavior core facility. For TubA experiments, daily injections were continued during the behavioral testing period. To minimize interference, injections were performed in the afternoon after the completion of each test. Before each test, mice were habituated to the room for at least 30 min and tests were performed under normal light conditions. Mice were handled once a day for 3 days prior to the first test. Muscle force of each animal was measured using a Grip Strength Meter (Chatillion DFE II, Columbus Instruments) with all paws (hindlimb grip strength test). The mouse was moved closer to the meter until it had a firm grip on the probe. The mouse was pulled horizontally away from the bar at a speed of ∼2.5 cm/s, until it released the probe. The value of the maximal peak force was recorded (gF). This was repeated five times for each animal, with a waiting time of 10 – 15 s between each measurement. In each animals at 30 day, the best score was defined as the specific maximal force. The grip strength measurements were conducted by the same investigator in order to limit variability and were performed in a random order. The investigator performing the measurements was blinded as to the treatment group of each individual mouse upon testing.

### Cell culture

C2C12 cells (ATCC) were seeded on matrigel-coated (matrigel® matrix, Corning) 35 mm-diameter plates and were maintained as myoblasts in Dulbecco’s modified Eagle medium (DMEM) supplemented with 10% fetal bovine serum and 1% penicillin-streptomycin (Multicell). Then cells were differentiated in differentiation medium (DMEM medium supplemented with 2% horse serum, Bio Media Canada). Cells grown in 35 mm diameter plates were treated for either Western blot or immunofluorescence. For Western blot, cells were collected by trypsinization, washed with PBS, centrifuged, and stored at -20°C until used. For immunofluorescence, cells were fixed for 20 min in PBS-4% paraformaldehyde at room temperature, washed in PBS and stored at 4°C until used.

### Drug, antibodies, and other reagents

C2C12 cells were treated with different drugs: tubastatin A (TubA at 5 µM, APExBIO, #A4101), tubacin (TBC, at 5 µM, Sigma, #SML0065) and SB 431542 (SB43, at 10 µM, Tocris, # 1614). Recombinant Human TGF-β1 (rhTGF-β1, at 10 ng/mL, # 100-21) was purchased from PeproTech. All primary antibodies used in this study are presented in table 1. Secondary antibodies used for immunofluorescence studies were coupled to Alexa-Fluor 488 or Alexa-Fluor 546 (Molecular Probes); or to Cy3 or Cy5 (Jackson ImmunoResearch Laboratories). Secondary antibodies used for Western blotting were either horseradish peroxidase (HRP)-coupled anti-rabbit-IgG polyclonal antibodies (Jackson ImmunoResearch Laboratories) or HRP goat anti-mouse-IgG antibodies (Millipore). To visualize NMJ for immunofluorescence studies, we used α-Bungarotoxin at 5 µg/mL conjugates with either Alexa-Fluor 488 (Molecular Probes) and DAPI (D9542; Sigma-Aldrich) was used to stain nuclear DNA. To visualize and quantify proteins on Western blot, we used 2,2,2-Trichloroethanol^110^ (TCE, Sigma, #T54801).

### Preparation of muscle and C2C12 cells homogenates

TA muscles were collected from adult mouse hindlimbs and dissected muscles were crushed on dry ice. Muscle powder resuspended in urea/thiourea buffer [7 M urea, 2 M thiourea, 65 mM chaps, 100 mM DTT, 10 U DNase I, protease inhibitors (Complete; Roche/Sigma-Aldrich)] and protein concentration was determined using CB-X Protein Assay kit (G-Bioscience, St. Louis, MO). After trypsination, C2C12 cells were solubilized in RIPA buffer [50 mM Tris– HCl, pH 8.0, 150 mM NaCl, 1% NP-40, 0.5% sodium deoxycholate, 0.1% SDS and protease inhibitors (Complete; Roche/Sigma-Aldrich)]. Protein concentration was determined using the BCA protein assay kit (Pierce/ThermoFisher Scientific) as per the manufacturer’s recommendations.

### Western blot

Five to twenty µg of total proteins were separated by SDS-PAGE supplemented with 0.5% TCE and transferred onto nitrocellulose or PVDF membranes. Non-specific binding was blocked with 4% bovine serum albumin (BSA, Euromedex) diluted in 1X PBS supplemented with Tween 0.1%, and membranes were incubated with primary antibodies. After thorough washing with 0.1% Tween 1X PBS, membranes were incubated with horseradish peroxidase (HRP)-conjugated secondary antibodies (Jackson Immunoresearch Laboratories/Cederlane). After additional washes, signals were revealed using ECL substrate reagents (Bio-Rad) and acquisitions were done using a ChemiDoc^™^ MP Imaging Systems (Bio-Rad) or autoradiographed with X-Ray films (Fisher Scientific). Quantifications based on TCE membrane were performed with the Image Lab software (Bio-Rad) or FIJI software (ImageJ 2.0.0-rc-69/1.52n, National Institutes of Health, Bethesda, MD).

### Muscle histology and histochemistry

Frozen TA muscle samples were placed at -20 °C into the cryostat (HM 525 NX, Thermo Fisher Scientific) for at least 20 min before further processing. 10 µm thick TA muscle sections were transversally cut. TA muscle cross-sections were stained with Hematoxylin and Eosin dyes. Sections were dehydrated using 70%, 90%, and 100% ethanol solutions and washed with toluene. The sections were mounted using Permount (Fisher Scientific) and visualized using an epifluorescent EVOS FLAuto2 inverted microscope. Percentage of central nucleation was determined by counting the total number of muscle fibers and the number of centrally nucleated muscle fibers from 6 to 8 cross-sectional views using the Northern Eclipse Software (NES, EMPIX Imaging).

### Immunofluorescence microscopy and image acquisition

Incubations with primary antibodies in PBS-0.1% Tween 20 were performed either at room temperature for 60 min (C2C12 cells) or at 4°C overnight (isolated dissociated muscle fibers) and washed. After incubation for 1-3 hours at room temperature with fluorescent secondary antibodies, DNA nuclear were stained with DAPI for 10 min. Coverslips were mounted on microscope slides with FluorSave^™^ reagent (Calbiochem). Images were captured at RT on either a Zeiss LSM880 microscope (Carl Zeiss) with an AiryScan1 detector equipped with a 63× 1.4-NA objective at INMG or a Zeiss Axio Imager M2 (Carl Zeiss) upright microscope equipped with either a 63× 1.4-NA or a 10× 0.45 NA objectives and AxioCam mRm CCD detector at the University of Ottawa Cell Biology and Image Acquisition core facility. All images were processed with either the ZEN blue software, Zeiss AxioVision software (Zeiss, Oberkochen, Germany), Photoshop CS5 (Abobe Systems, San Jose, CA, USA) or FIJI software (ImageJ 2.0.0-rc-69/1.52n, National Institutes of Health, Bethesda, MD). Images were analyzed in a blinded manner by randomly renaming file names with numbers using the ‘name_randomizer’ macro in ImageJ^77^.

### Quantitative Microtubule network lattice analysis

Using ImageJ, microtubule organization was visualized with vertical (yellow bars) and horizontal (blue bars) line scans. Using a recently developed directionality analysis program^78^ (TeDT), microtubule network lattice directionality was calculated for all mouse lines. A two-way ANOVA was used to assess the effect of microtubule intersection angle across groups. two-sided U test (Mann-Whitney) post hoc measures were used to determine the extent of differences between groups. Significance was set at P < 0.05.

### Quantitative analysis of compactness and fragmentation index by ‘NMJ-morph’ methodology

For accurate analysis, each image was captured a single en-face NMJ. NMJs that were partially oblique to the field of view were only included if the oblique portion constituted less than approximately 10% of the total area. To quantify compactness and fragmentation index, images were analyzed thanks to ‘NMJ-morph’ methodology^111^. The compactness of AChRs at the endplate was defined as follows: Compactness = ((AChR area)/ (endplate area)) ×100. Fragmentation index was calculated whereby a solid plaque-like endplate has an index of (0), and highly fragmented endplate has an index that tends towards a numerical value of (1): Fragmentation index = 1-(1/ (number of AChR cluster)). The basic dimensions of the post-synaptic motor endplate were measured using standard ImageJ functions. ‘NMJ-morph’ is used to quantify the number of discrete AChR clusters comprising the motor endplate.

### Statistical analyses

All statistical analyses were performed using Prism 6.0 (GraphPad Software, La Jolla, USA). Data are given as mean ± SEM. Student’s t-test was used if datasets belong to a normally distributed population with an n > 30. Otherwise, the nonparametric, two-sided U test (Mann-Whitney) was applied. Data distribution was assumed to be normal, but this was not formally tested. For a multiple factorial analysis of variance, two-way ANOVA was applied. P-values under 0.05 were considered statistically significant (shown as a single asterisk in figures); p-values under 0.01 were considered highly statistically significant (shown as two asterisks in figures); p-values under 0.001 were considered very highly statistically significant (shown as three asterisks in figures).

## Acknowledgements

We thank John Lunde, Jean Luc Thomas, Amanda Tran, and Laurent Coudert for expert technical help and fruitful discussion.

## Author contributions

A.O, B.J.J and L.S conceived the study, designed the project, and obtained grant funding. A.O and A.R.C performed immunofluorescence and all animal experiments. A.O and I.S performed TGF-β experiments. A.O and Y.G.G. performed mTOR experiments. A.O. and P.L performed all the Western blots. A.O performed all the culture cell experiments. A.O and L.S wrote the first draft manuscript. A.O, A.R.C, I.S, Y.G.G, V.M, R.M, P.L, B.J.J and L.S analyzed, interpreted the data, reviewed, finalized the manuscript, and provided comments and edits.

## Additional information

Supplementary information is showed in Table 1: classification of antibodies.

