## Supplemental Table 1 for "Pharmacological inhibition of HDAC6 downregulates TGF-β via Smad2/3 acetylation and improves dystrophin-deficient muscles"

### Table 1: Antibodies

| antibody name | type | dilution | | provider |
|---|---|---|---|---|
| | | Immunostaining | Western blotting | |
| β-tubulin | mouse monoclonal | 1:500 | | Sigma-Aldrich, clone TUB2.1, #T5201 |
| acetylated tubulin | mouse monoclonal | 1:200 | 1:2,000 | Sigma-Aldrich, clone 6-11 B-1 #T7451 |
| α-tubulin | mouse monoclonal | | 1:1,000 | Sigma-Aldrich, clone B-5-1-2, #T6074 |
| Histone H3 (D1H2) | rabbit monoclonal | | 1:1,000 | Cell Signaling Technology, #4499 |
| acetyl-Histone H3 (Lys9) | rabbit polyclonal | | 1:1,000 | Sigma-Aldrich, #07-352 |
| Utrophin A | mouse monoclonal | 1:200 | 1:500 | Novocastra, Leica biosystems, #NCL-DRP2 |
| β-dystroglycan | mouse monoclonal | 1:400 | 1:500 | Novocastra, Leica biosystems, #NCL-b-DG |
| Atrogin-1 (MAFbx) | rabbit polyclonal | | 1:1,000 | ECM Biosciences, #AP2041 |
| MuRF1 | mouse monoclonal | | 1:1,000 | Abcam, #ab57865 |
| Col1A1 | goat polyclonal | | 1:200 | Santa Cruz Biotechnology, #sc-25974 |
| CTGF | rabbit polyclonal | | 1:1,000 | Abcam, #ab6992 |
| TSC1 | rabbit polyclonal | | 1:1,000 | Cell Signaling Technology, #4906 |
| mTOR | rabbit polyclonal | | 1:1,000 | Cell Signaling Technology, #2972 |
| Phospho-p70 S6 Kinase (Thr389) | rabbit polyclonal | | 1:1,000 | Cell Signaling Technology, #9205 |
| p70 S6 Kinase | rabbit polyclonal | | 1:1,000 | Cell Signaling Technology, #9202 |
| Phospho-S6 Ribosomal Protein (Ser240/244) | rabbit polyclonal | | 1:1,000 | Cell Signaling Technology, #2215 |
| Phospho-S6 Ribosomal Protein (Ser235/236) (D57.2.2E) | rabbit monoclonal | | 1:1,000 | Cell Signaling Technology, #4858 |
| S6 Ribosomal Protein (5G10) | rabbit monoclonal | | 1:1,000 | Cell Signaling Technology, #2217 |
| Phospho-4E-BP1 (Thr37/46) (236B4) | rabbit monoclonal | | 1:1,000 | Cell Signaling Technology, #2855 |
| Phospho-4E-BP1 (Thr70) | rabbit polyclonal | | 1:1,000 | Cell Signaling Technology, #9455 |
| 4E-BP1 | rabbit polyclonal | | 1:1,000 | Cell Signaling Technology, #9452 |
| Phospho-Smad2 (Ser465/467) /Smad3 (Ser423/425) (D27F4) | rabbit monoclonal | | 1:1,000 | Cell Signaling Technology, #8828 |
| Smad2/3 (D7G7) | rabbit monoclonal | 1:400 | 1:1,000 | Cell Signaling Technology, #8685 |
| Acetyl-SMAD2/SMAD3 (Lys19) | rabbit polyclonal | | 1:1,000 | Invitrogen, #PA5-76015 |
| Smad3 (phospho S423 + S425) | rabbit polyclonal | 1:200 | 1:1,000 | Abcam, #ab52903 |
| laminin | rabbit polyclonal | 1:200 | | Sigma-Aldrich, #L9393 |
| GAPDH (HRP) | goat polyclonal | | 1:20,000 | Abcam, #ab85760 |
