## Supplementary figures and images for "Pharmacological inhibition of HDAC6 downregulates TGF-β via Smad2/3 acetylation and improves dystrophin-deficient muscles"

### Extended Data 2

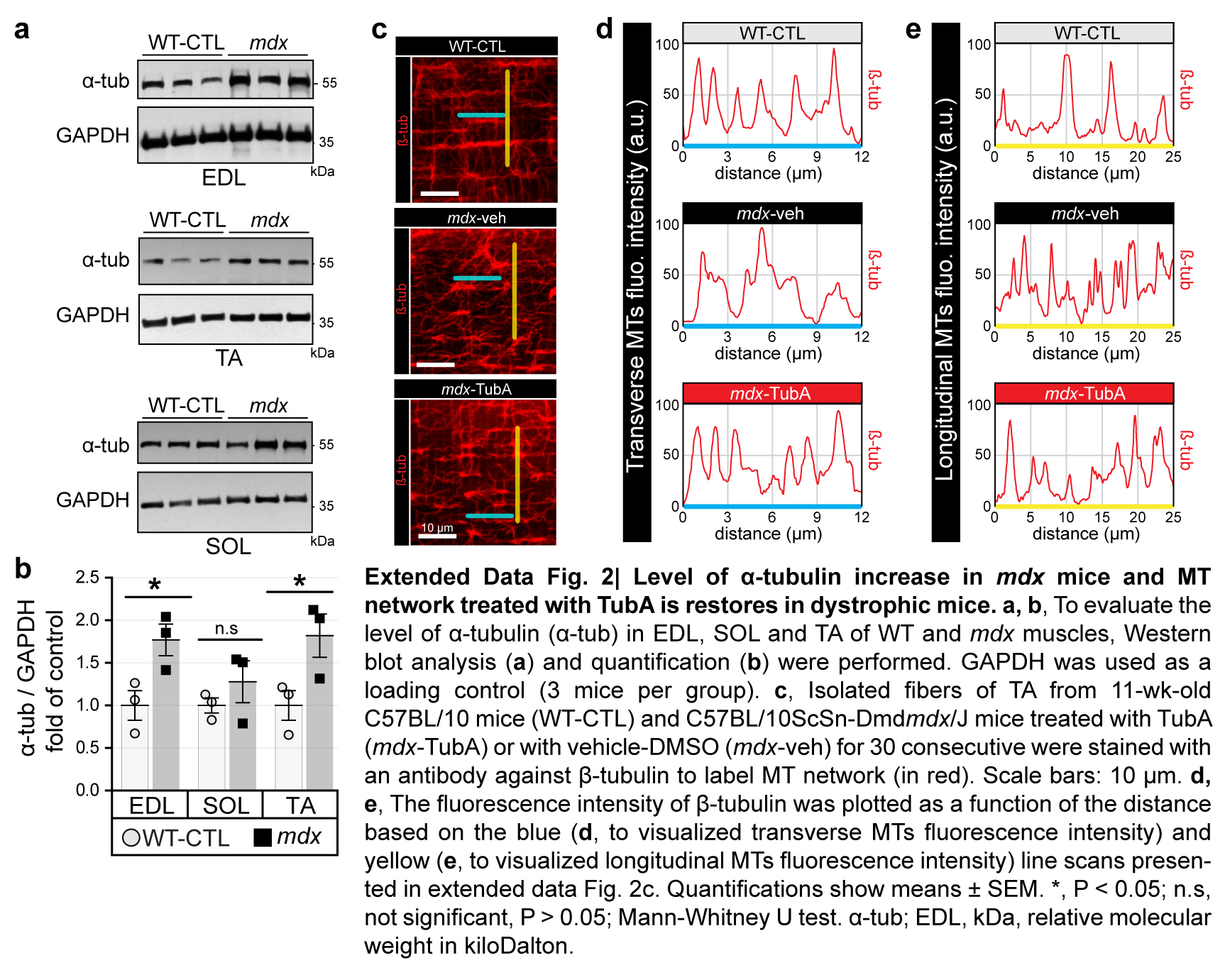

### Extented Data 1

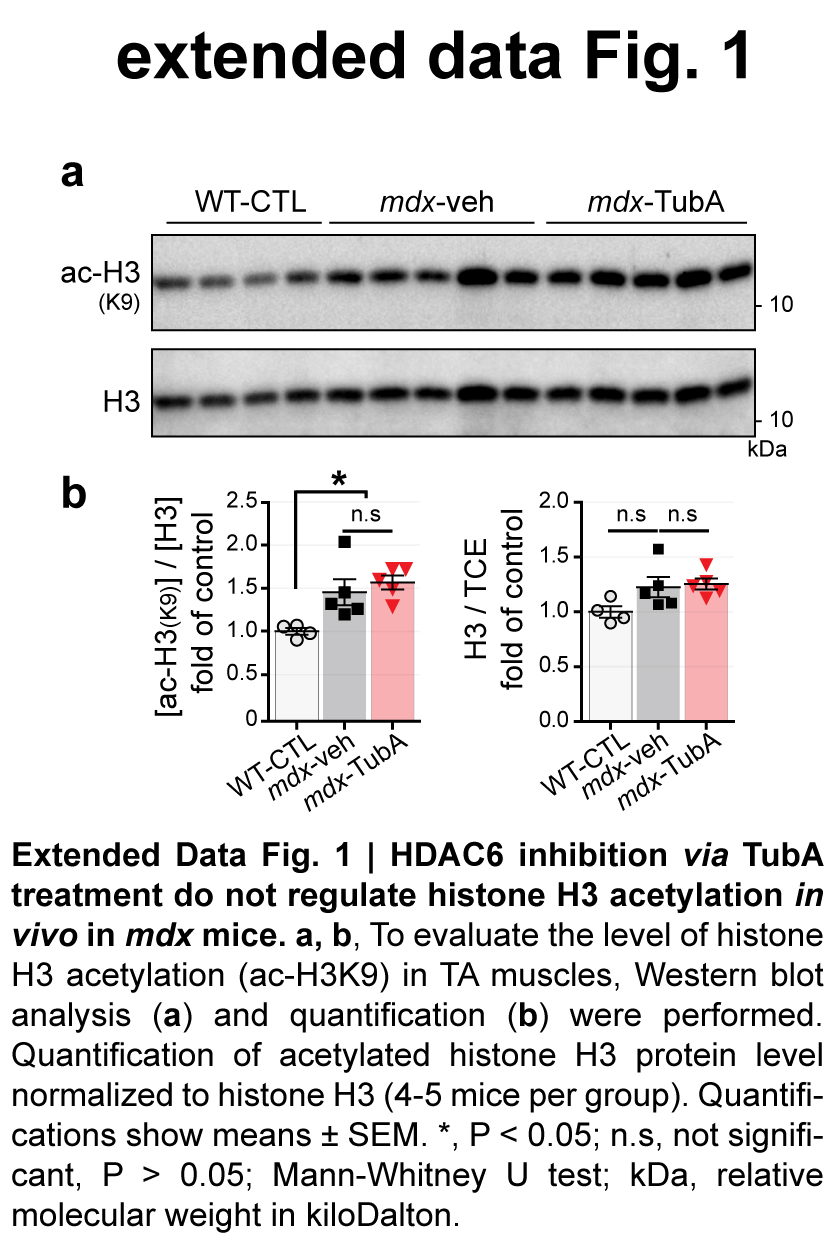

### Extented Data 3

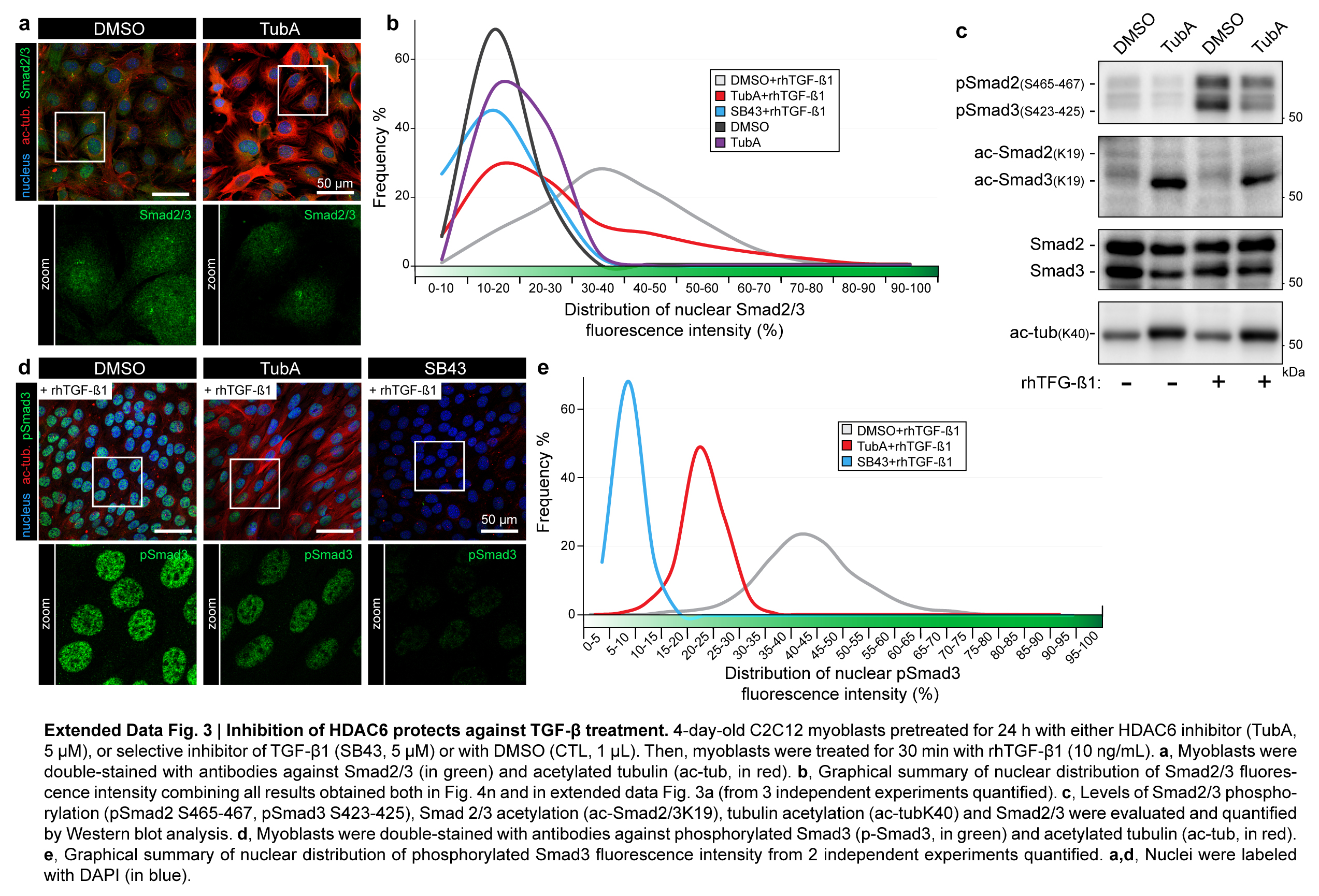
